# Increased Functional Connectivity Characterises the Prodromal Phase of the Migraine Cycle

**DOI:** 10.1101/2021.10.18.464798

**Authors:** Anne Stankewitz, Astrid Mayr, Enrico Schulz

## Abstract

**Introduction:** Episodic migraine is reflected by cyclic changes in behaviour and cortical processing. We aimed to identify how Functional connectivities change over the entire migraine cycle.

**Methods:** By using longitudinal neuroimaging and a whole-brain connectivity analysis approach, we tested 12 episodic migraine patients across 82 FMRI recordings during spontaneous headache attacks with Follow-up measurements over the pain-free interval without any external stimulation.

**Results:** We Found that Functional connectivities linearly increased over the interictal interval. In the prodromal phase, we observed the strongest connections between the anterior agranular insula and the posterior orbitofrontal cortex with sensory, motor, and cingulate areas. The strengths of the connections dropped during the headache. Peak connectivity during the prodromal phase and its collapse during the headache can be regarded as a mechanism of normalising cortical processing.

**Conclusions:** The strongest connections during the ictal phase of the migraine cycle may contribute to the variety of symptoms of migraine attacks including headache, sensory hypersensitivity, and autonomous symptoms. We speculate about a malfunction at the molecular level in agranular frontal and insular regions, which needs to be addressed in subsequent studies.

**Article highlights:**

- We investigated functional connectivities over an entire migraine cycle.
- We found cycle-related connectivity changes for two proximate agranular regions.
- The prodromal increase and the collapse of connectivity during the headache may reflect normalising cortical processing.

## Introduction

Episodic migraine is a cyclic disease with recurring headache episodes, which are often the most disabling symptom. However, migraine attacks are more than just a headache; attacks typically last up to 72 hours and contain up to four phases with characteristic symptoms separated by interictal intervals with a variable length: the prodromal (premonitory), aura, headache, and postdrome phase ^1^.

### The cyclic nature of migraine disease

Evidence for cyclic changes in the migraineurs’ brain originates from early neurophysiological studies: EEG results revealed a lack of habituation to repetitive sensory stimulation ^2^ during the pain-free interval that normalised just before or during the headache attack ^3^. Migraine phase-dependent BOLD activity in the trigeminal nuclei ^4^ and the hypothalamus ^5,6^ has further been observed in fMRI studies. In addition, enhanced functional connectivities (FC) were observed between limbic regions during the preictal phase and between the pons and the hypothalamus during the headache attack ^7^. During the headache attack, FC alterations have been reported in pontine regions ^8^ and the thalamus ^9^. Recently, our group observed cyclic changes of sensory, thalamic and sensory networks: the activity within these networks linearly increased over the pain-free interval, with peak activity during the prodromal phase, but decoupling of network regions during the acute headache attack ^10^.

### Hypothalamus as a rhythmic generator of migraine attacks

The cyclicity of episodic attacks in migraineurs has been attributed to the hypothalamus due to its role in the regulation of biological rhythms and in maintaining internal homeostasis by controlling the endocrine and the autonomic nervous system ^11^. Several migraine symptoms and trigger factors of attacks are related to the hypothalamus ^5,12,13^. Hypothalamic alterations were indeed observed not only during the headache attack ^14^ but also prior to the headache during the prodromal phase ^5,6,15^.

### The importance of the preictal migraine phase

Previous research ^3,5,6,15^ as well as clinical features ^11^ of the disease (e.g. attack trigger and premonitory symptoms) suggest that the initiation of headache attacks most likely starts during the preictal - pain-free - interval, which may be hours or even days before the headache occurs. However, most imaging studies on migraine were utilising cross-sectional designs; longitudinal within-subject designs, which are necessary to shed light on the cyclicity of migraine-related mechanisms in the brain, are rare ^5,6,16^. In addition, connectivity studies that investigated patients at different migraine phases used *a priori* defined seed regions instead of taking the whole-brain connectivity into account^7,17^.

To expand our knowledge about the naturally developing and untriggered cyclic mechanisms of the migraineurs’ brain, we aimed to explore the trajectory of whole-brain FC during resting state-fMRI (rs-FMRI) over the entire migraine cycle. *First*, based on the clinical picture of the disease and previous imaging findings, we hypothesised hypothalamic alterations in connectivity along the migraine cycle. *Second*, due to sensory dysfunctions reported by migraine patients during and outside attacks, we hypothesised cyclic dysfunctions in functional connections involving sensory brain areas.

## Methods

### Subjects

Participants were recruited through word-of-mouth and by an email distributed to employees of the university hospital “Klinikum rechts der Isar”. 12 patients with episodic migraine (11 females and one male; 28±5 years) were included in the study (Supplementary Table 1). The time points of the attacks were equally distributed along the menstrual cycles of the female patients. Migraine diagnosis was given by a headache specialist and was based upon the classification of the International Headache Society ^18^. The patients did not report any other neurological or psychiatric disorders, were not taking preventative medication for migraine for at least six months, but were allowed to take their regular acute migraine medication *after* the recording of the headache attack (non-steroidal anti-inflammatory drugs or triptans). The interval between the intake of acute medication and the first postdrome recording was between 12h and 24h for each participant. Due to their effects on cortical perfusion, patients were not permitted to consume coffee or other caffeinated beverages 12h before the recordings. All patients gave their written, informed consent. The study was conducted according to the Declaration of Helsinki and approved by the Ethics Committee of the Technische Universität München (project number 90/14). All patients were remunerated for participation.

### Study Design

Migraine patients were tested repeatedly over an entire migraine cycle (Figure 1). The imaging time series for each patient started with the recording of a spontaneous and untreated headache attack within the first 6 hours after the beginning of the headache. We only recorded headache attacks that were reported with an intensity of moderate to severe, predefined as a minimum of “4” on a numerical rating scale ^19^ with the endpoints “0” (no pain) and “10” (extremely intense pain). Brain data were then recorded every 1-4 days (1.7±0.8 days; recordings: min=5, max=10; n=6.8±1.5) until patients informed us by phone about the following headache attack (which was not scanned) and the time series was completed with the last attack-free recording.

**Fig. 1.**
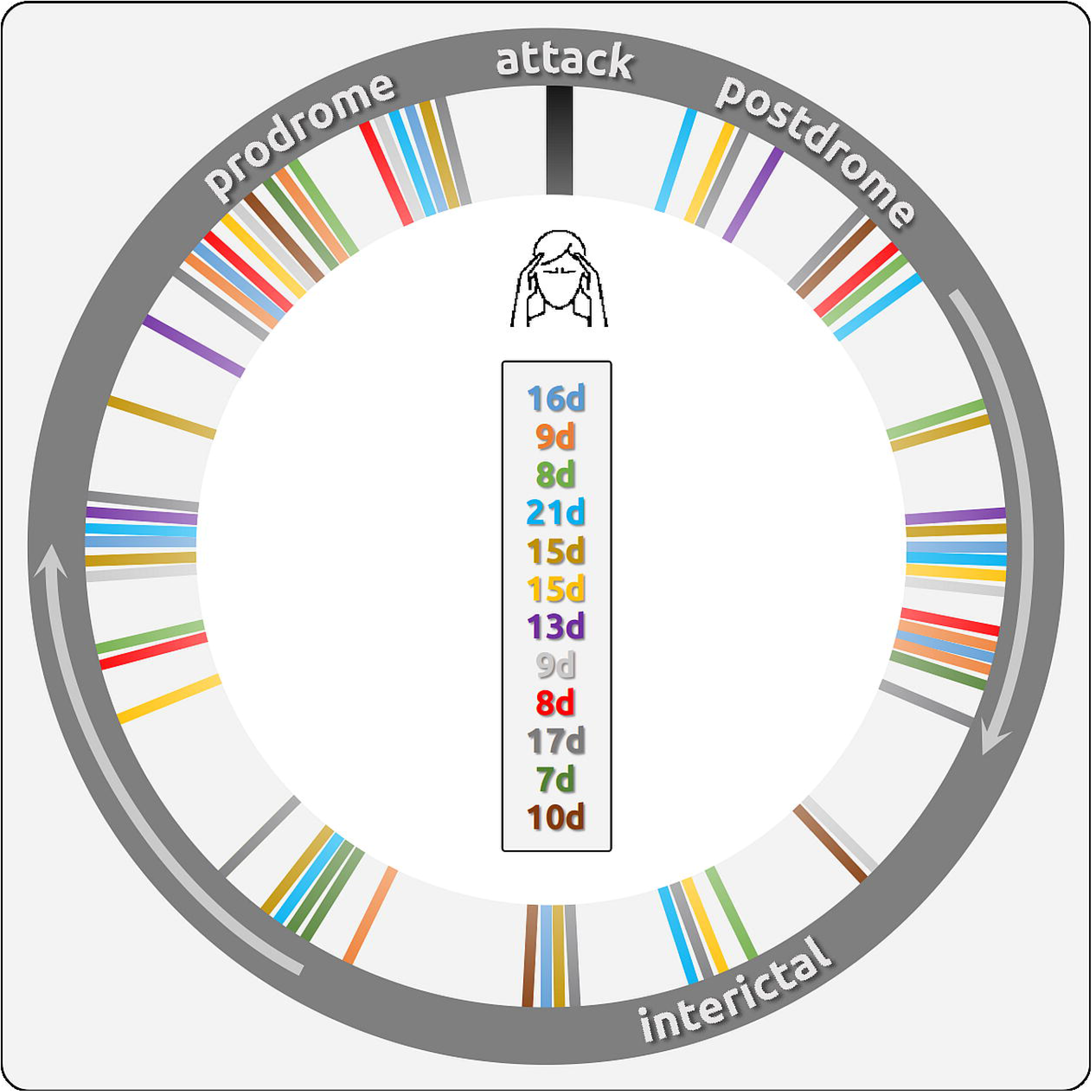
Circular time course of individual recordings between two migraine attacks. The figure integrated the recording days into the circular representation of a migraine cycle. Each colourful line indicates a separate recording. Same colours belong to the same patient. The number of days between the first recorded headache attack and the subsequent headache attack is shown in the middle and indicates the different lengths of each patient’s migraine cycle. We recorded one migraine cycle per patient.

### Image Acquisition

MRI data were collected on a 3 tesla scanner (Ingenia, Philips, The Netherlands) using a 32-channel head coil. Patients were instructed to remain awake and relaxed with their eyes closed. For the 300 volumes of resting-state data, we used the following parameters: TR=2000 ms; TE=30 ms; FOV=192 × 192 mm^2^; flip angle=90°; number of slices=37; voxel size=3 × 3 x 3 mm^3^ (0.29 mm gap). For image registration, a high resolution T1-weighted anatomical image was collected with: TR=9000 ms, TE=4 ms, flip angle=8°, FOV=240 x 240 x 170 mm^3^; number of slices=170; voxel size=1.0 x 1.0 × 1.0 mm^3^). Field maps were acquired in each session to control for B0-effects; 64 slices, TR=960 ms, FOV=192 × 192 mm^2^; voxel size=2.0 x 2.0 × 2.0 mm^3^, 0.2 mm gap between slices. TE=6 ms / 10.55 ms, flip angle 60°.

### Image Preprocessing

The data were preprocessed with FSL ^20^. The Melodic toolbox was used to execute brain extraction, high-pass filtering with a frequency cutoff of 1/100 Hz, spatial registration to the MNI template, corrections for head motion during scanning, and spatial smoothing (5mm FWHM). A distortion correction of the images was used based on field maps. The data were further semi-automatically cleaned of artefacts with an independent component analysis (ICA) through Melodic ^21^. The number of components had been automatically estimated by the software and artefact-related components were removed from the data. Head movement during scanning did not exceed 2mm or 2° in any direction.

The time series of functional volumes were projected to surface space by using the “Connectome Workbench” package. Regions of interest (ROIs) were defined by subdividing the cortical surface into 180 regions per hemisphere ^22^. 65 further regions including the cerebellum and subcortical regions, such as the periaqueductal grey (PAG), the thalamus, cerebellar subregions, and the amygdala, were also included. 21 of the subcortical regions are included in the Glasser template. The cerebellar and further subcortical regions were derived from the FSL atlas. The remaining ROIs were created based on the literature, e.g. the periaqueductal grey. The time courses for all voxels of cortical activity for a specific region of the Glasser Atlas, e.g. the middle insula, were extracted. Nuisance variables (McFLIRT motion parameters, their squares and temporal derivatives with squares; a total of 24 regressors), as well as outliers, were regressed out from the data. Outliers in the fMRI data are defined by the framewise displacement (FD) and DVARS. A volume is defined as an outlier if it exceeds one of the following thresholds: FD ≧ 0.2mm or DVARS ≧ (the 75th percentile + 1.5 times the InterQuartile Range). Outliers are marked for each subject and then included in the regression as a vector of zeros (non-outlier) and ones (outlier). We computed principal component analyses (PCA) separately for each ROI and subject and selected the first component (Version R2018a, Mathworks, USA). Time courses of the 425 components were correlated using Kendall’s τ coefficient.

### Statistical Analyses

We explored the cyclic change of cortical resting-state connectivity between the 425 regions over the migraine interval. To investigate how the cortical connectivities evolve over the trajectory of measurements, we computed linear mixed-effects models (LME) for each pair of brain regions and related the time points within the migraine cycle to the strength of connectivity quantified by Kendall’s τ. We created a time vector for each patient’s migraine cycle and encoded the day of the recording by assigning numbers between 1 and 2. The numbers 1 or 2 were assigned to the measurement during the headache attack, depending on the following two different trajectories of map strengths (Figure 2) in the brain ^5^.

**Fig. 2.**
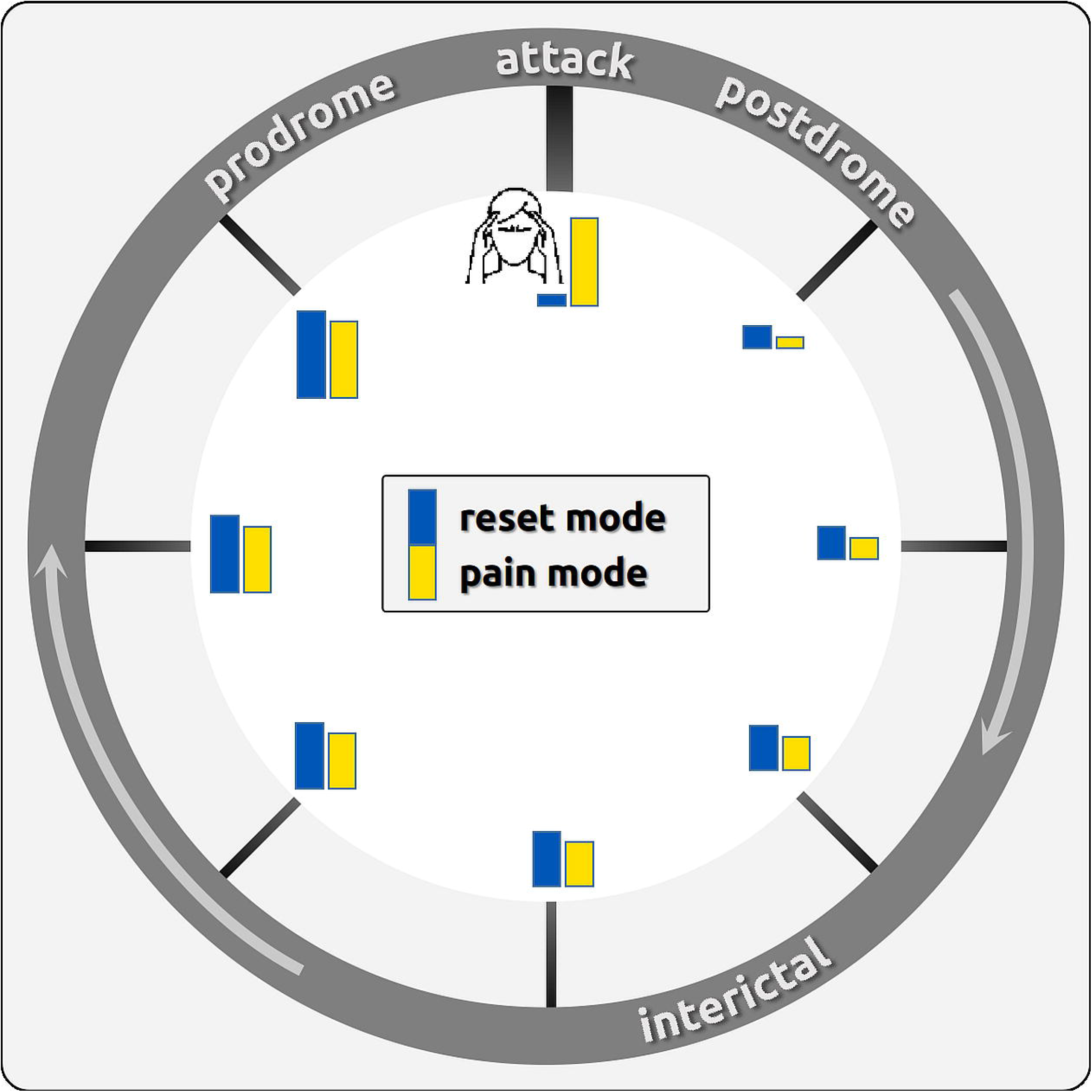
Circular time series of migraine-related brain processes. Two hypothetical time series (trajectories) of migraine-related brain processes were modelled in the statistical analysis. In the first time series (blue) the brain processes drop during the headache attacks; the brain processes would be “reset” during attacks, then would resemble the connectivity on the day after the attacks. In the second time series (yellow), the brain processes would reach their maximum during the attacks and are similar to the days before attacks. These processes could be used as a biomarker for an impending migraine attack.

> **Trajectory 1 (“reset mode”)**: In this model, we aimed to detect the resting-state connectivity that has its peak (positive or negative) just *before* the headache attack starts and then drops to the *level (positive or negative) during the headache*. From here, the map strength linearly increases (or decreases) over the migraine cycle to the next attack. In this hypothetical time course, the map strength on the first day *after the attack* would be similar to the brain connectivity during the attack. This trajectory can be interpreted as a cortical “reset” mechanism and is in line with neurophysiological studies reporting a habituation deficit in pain-free migraineurs that normalise during headache attacks ^3^. For example, for 5 measurements over 10 days, the following vector is used to reflect the trajectory 1: *1 (=attack), 1.2, 1.4, 1.6, 1.8*.

> **Trajectory 2 (“pain mode”)**: In this model, we aimed to detect the resting-state connectivity that has a peak *during* the headache attack and drops to the level (positive or negative) during the postdrome phase. From there, we assume a linear increase (or decrease) over the migraine cycle towards the next attack. We hypothesise increased brain connectivity in regions that contribute to the processing of migraine symptoms, e.g. pain, increased sensitivity to light, sound, and odours, and vegetative complaints. In this hypothetical time course, the brain connectivity on the day *prior to the attack* would be similar to the brain connectivity during the headache attack and is suggested to reflect the steadily increasing excitability of the brain. Similar to the above-mentioned example with 5 measurements over 10 days, the following vector is used for trajectory 2: *2 (=attack), 1.2, 1.4, 1.6, 1.8*.

The statistical analysis for the trajectory of resting-state connectivity has been performed in Matlab. To explore the relationship between the fluctuating map strength and the dependence of the time point of the migraine cycle, we computed LMEs that related the longitudinal recordings of resting-state connectivity to the number vector of recording days:

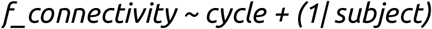

The included fixed effect (f_connectivity ~ cycle) essentially describes the magnitudes of the population common intercept and the population common slope for the dependency of cortical data on the factor time. The added random effect (e.g. 1 | subject) is used to model the specific intercept differences for each subject. In other words, the model estimates the relationship between the cortical processes in dependence on their occurrence in the migraine cycle (fixed effect). Please note that the statistical terms of fixed effect and random effect are differently used in the common neuroimaging software package SPM as compared with the standard nomenclature in statistics that we are following here. T-values are computed voxel-wise as quotients between the beta estimates and the standard errors of the equation.

Based on 5000 randomisations (see Supplementary Materials), the statistical threshold (p<0.05, equivalent to t>5.195) was determined using the “palm_datapval.m” function publicly available in PALM ^23^. This approach corrects for multiple testing. Connectivity figures were generated using NeuroMArVL ([link](https://immersive.erc.monash.edu/neuromarvl/?save=45220d30-ce3b-4e17-922c-88964a883beb_31.16.56.20)).

## Results

To determine the cyclic changes of the functional connectivity in the migraineurs’ brain between two migraine attacks, we computed linear mixed-effects models (LME) and analysed how the resting-state functional connectivity is evolving depending upon the time point within the migraine cycle. All 66 significant pairs of connectivity for trajectory 1 have a positive mean across all 81 Kendall’s τ values and 94% of these pairs (n=62) show a higher mean than standard deviation, indicating that the effects are predominantly based on a stronger co-variation of the time courses between two brain regions in the prodrome phase compared to a declined connectivity during the headache attack as reflected by Kendall’s τ coefficients around 0. The discussion of the results is based only on these interpretable 62 pairs of brain regions.

The findings are presented in Figures 3a, 3b, and 3c. A confusion matrix gives a more global impression on the direction of connectivity changes throughout the migraine cycle (Figure 3a), a circular plot shows all significant connections (3b), and the best-connected brain regions are presented on a 3D brain (3c). All figures show different aspects of the same underlying results. Two possible time courses were explored:

**Fig. 3a.**
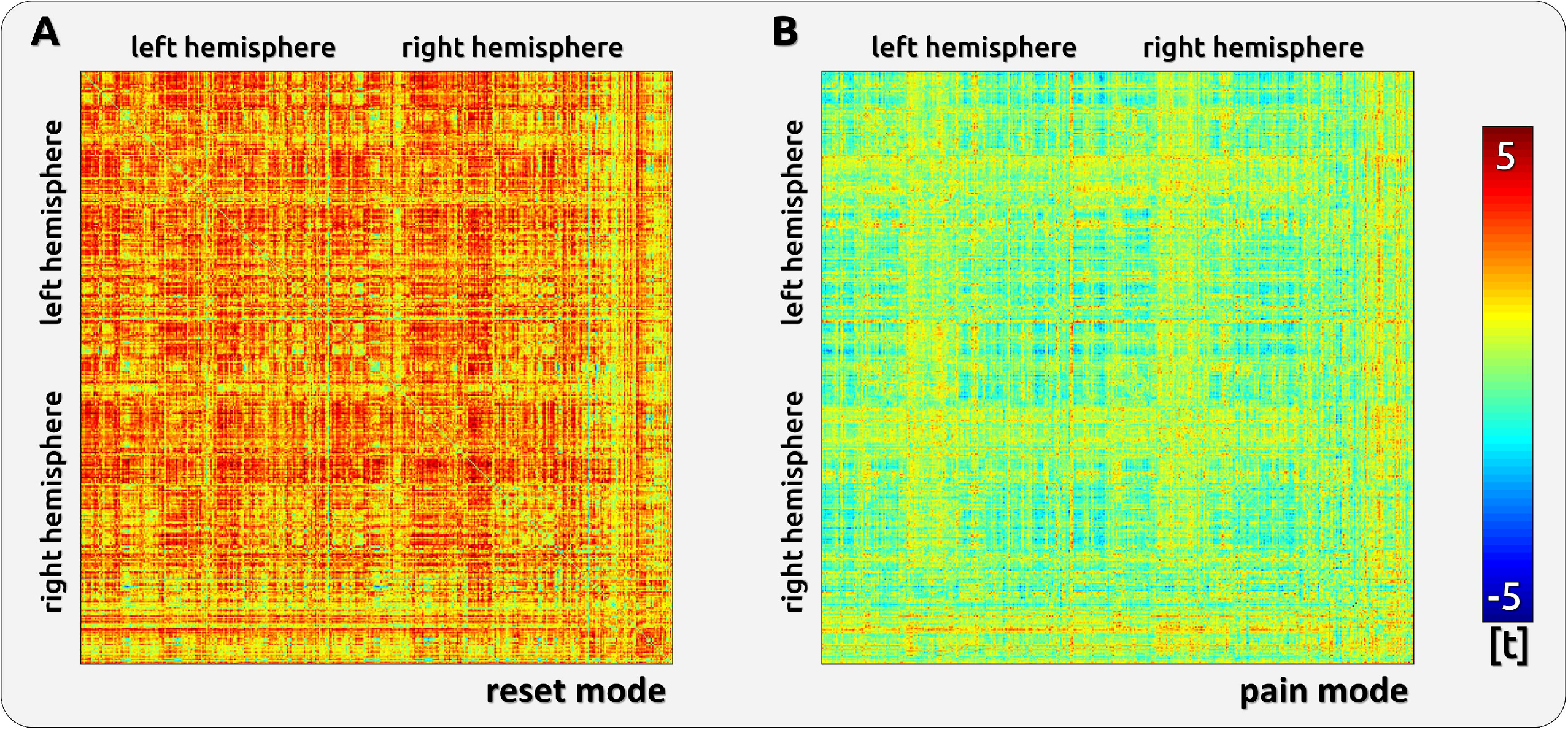
Confusion matrix on migraine cycle-related connectivity changes across all 425 brain regions. The left matrix (A) shows the data for trajectory 1 (reset mode) that has its peak at a pain-free time point just before the headache attack. All of these trajectories show an increase of connectivity over time with a drop during the headache attack. There was no decreasing connectivity towards a minimum at the time point just before the attack. The right matrix (B) has only a single significant pixel that represents the highest connectivity between the hypothalamus and the LC during a headache attack (pain mode).

**Fig. 3b.**
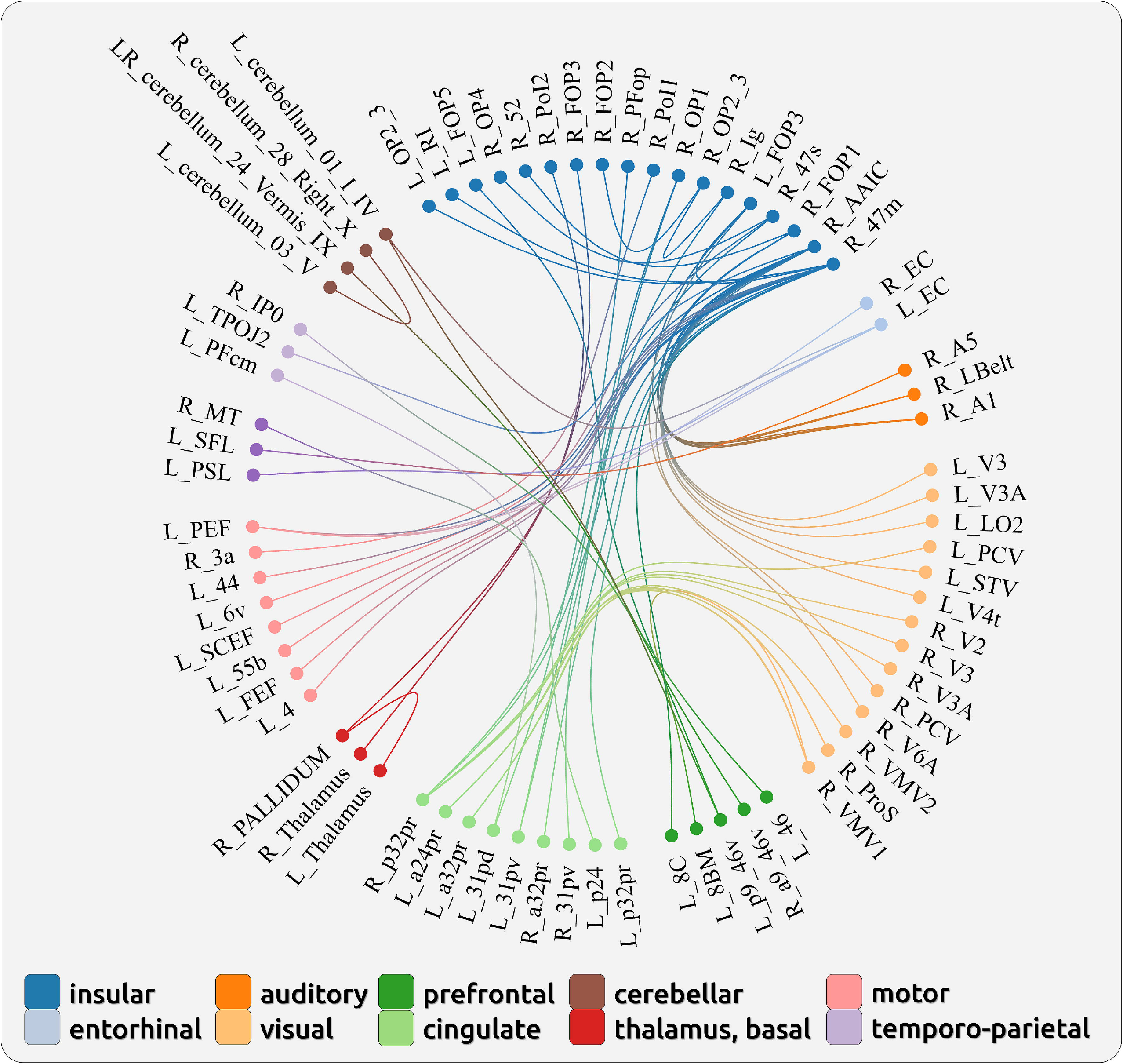
Cycle-related connectivity changes following the “reset mode” trajectory. The figure shows a circular plot ([link](https://immersive.erc.monash.edu/neuromarvl/?save=45220d30-ce3b-4e17-922c-88964a883beb_31.16.56.20)) representing all significant connections.

**Fig. 3c.**
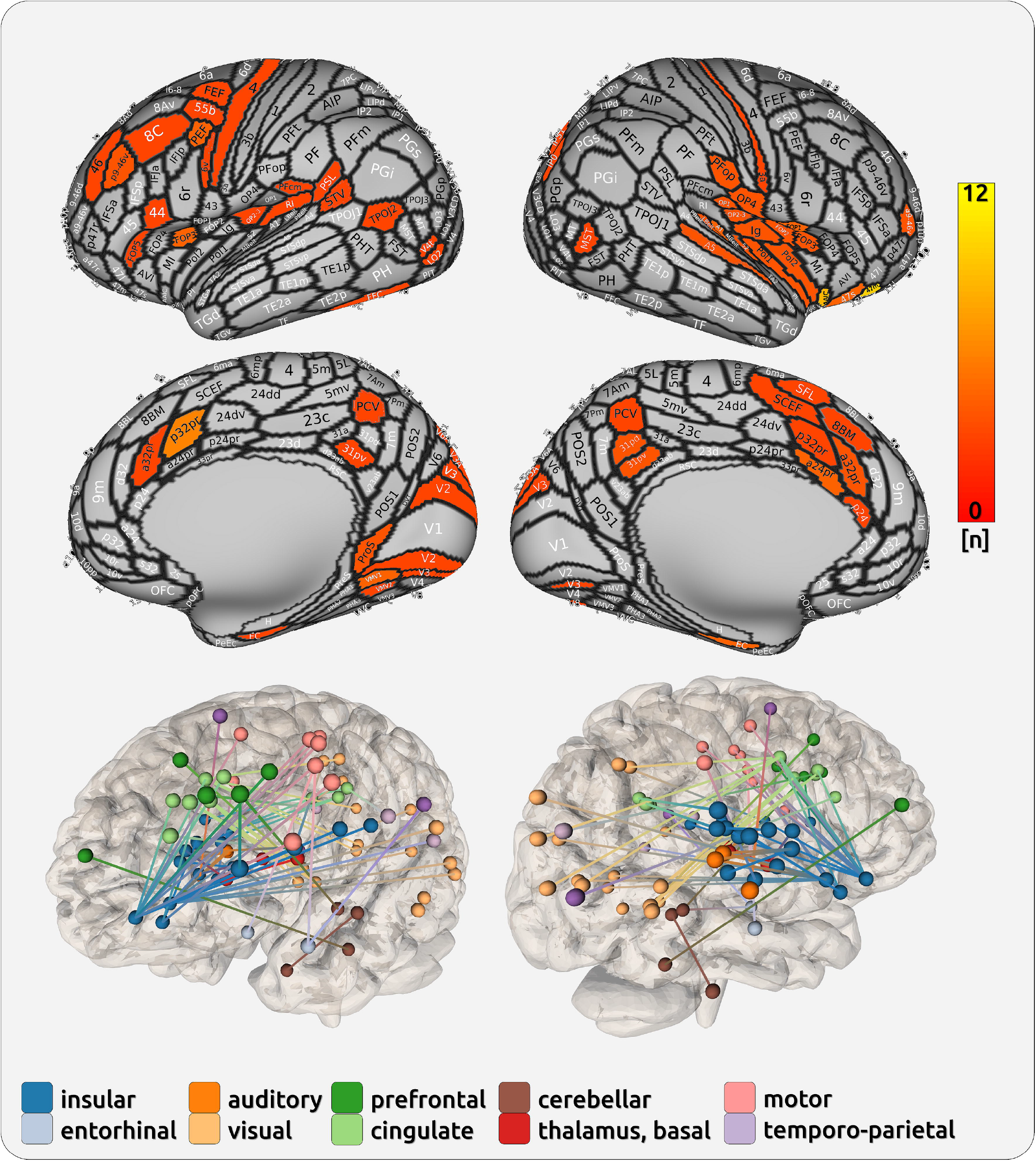
Cycle-related connectivity changes following the “reset mode” trajectory. The figure shows the connected brain regions. The colour coding in the upper part indicates the number of connections with the highest number for the area 47m (12) and the aAIC (10).

### Trajectory 1

For this trajectory, the functional connectivity between brain regions increased/decreased linearly over the interictal interval towards the attack, reached its peak/trough prior to the beginning of the headache, and dropped back to normal during the headache. We found three larger complexes of brain regions that account for most of the significant connections: the insular cortex and surrounding regions, the dorsal posterior cingulate cortex (PCC; Brodmann area 31), and the dorsal anterior cingulate cortex (ACC; Brodmann area 24 and 32).

#### Insula and adjacent interior frontal cortex

For the insular cortex and its surrounding sub-regions, the most affected region is the area 47m (12 connections), which reached its peak/trough prior to the beginning of the headache and drops its connection during the headache attack to insular and parietal opercular regions, the primary somatosensory cortex, regions that process eye movements, and the dorsal posterior cingulate cortex. The anterior agranular insular cortex (AAIC; 10 connections) exhibits a steady increase of connectivity prior to the attack and shows a drop of connectivity predominantly to areas related to sensory functions (auditory - lateral belt region, primary auditory cortex; visual - V3, V3A, V4t, V6A; visual integration areas - V6A, TPOJ, superior temporal visual area).

#### Posterior Cingulate Cortex

We found several connections between the PCC (BA 31) and the insular-opercular regions.

#### Anterior Cingulate Cortex

The dorsal ACC showed multiple connections to the inferior frontal regions (BA 47), visual (V1 to V3), and higher-order visual areas (prostriate and ventromedial visual areas 1 and 2).

### Trajectory 2

There are no significant effects for this model.

## Discussion

The present longitudinal fMRI study aimed to detect the whole-brain functional connectivity in episodic migraineurs over the migraine cycle. By taking the time point of the attack into account, we analysed two possible connectivity trajectories: (1) a linear trajectory (increase or decrease) towards the attack with peak connectivity during the prodromal phase, followed by connectivity “reset” to the baseline level during the headache (“reset mode”), or (2) a linear trajectory (increase or decrease) towards the attack with peak connectivity during the headache (“pain mode”). Significant results were found only for the first trajectory.

### Trajectory of the “reset mode”

The present study suggests that pathological processes are developing early on and can be observed and potentially treated before the painful phase of the headache attack is initiated. This is in line with previous neurophysiological research that has identified deviant sensory processing in the interictal phase: studies reported a lack of habituation to sensory stimulation during the interictal phase of the migraine cycle followed by a “normalisation” of habituation just before and during the headache attack ^3,24^. This migraine-cycle-dependent cortical processing has been interpreted in a way that the headache attack may represent a “reset-to-normal” of cortical processing, particularly in regard to sensory processes ^25^. These findings served as the theoretical basis for the generation of the vector, where we also modelled a similar cortical processing in the headache phase and the postdrome phase with a linear increase towards the prodrome phase. Here, we identified three major regions that may represent these deviances.

#### Connectivity of insular regions

There are a number of connections in subregions of the bilateral insular cortex affected throughout the migraine cycle. The best-connected insular region is the agranular anterior insular cortex (AAIC), which exhibits a steady increase of connectivity towards the attack and shows an ictal drop of connectivity particularly to sensory areas (auditory, higher-order visual). As the AAIC is also anatomically connected to olfactory areas, the region has been related to the processing and integration of sensory information ^26^ as well as part of the salience network ^27^. The anterior insula is also involved in the cognitive aspects of processing pain ^28^ and in maintaining working memory functions in mice ^29^ and men ^30^. This is in line with our previous work showing a similarly increasing connectivity from the hypothalamus to the right anterior insular cortex ^5^. We suggested that the steadily increasing hypothalamic-insular connectivity would serve as a control mechanism for the activity in insular sub-areas. However, this hypothalamic control may require a steadily increasing effort over the migraine cycle, probably due to the increasing connectivity from sensory areas to the anterior insula that we find in the present investigation.

#### Connectivity of inferior frontal region 47m

The best-connected region is the right area 47m, which is located in the inferior frontal cortex. An involvement of this region in the context of pain and migraine has been found in the seminal work of Denuelle and colleagues ^14^. Using PET during migraine attacks, the Brodmann Area 47 was active during the headache as well as after the successful treatment of the headache with analgesic medication, suggesting that this region is involved in the underlying migraine-related cortical processes.

In the present study, the area 47m exhibits a migraine-related cyclic trajectory predominantly with regions in the left insula and with the bilateral posterior part of the anterior cingulate cortex (BA 32). There is a vast amount of literature showing the involvement of these regions in the processing of pain ^31^ partly due to the increased but unspecific salience of the perceived pain ^32,33^. Our data showed peak connectivity between area 47m and BA 32 during the prodrome phase and lowest connectivity during the headache. We may see a functional distinction in the migraine-related impairments of sensory processing (i.e. the lack of habituation). While visual and auditory processes are directly bound to the AAIC, the area 47m also exhibits connections to the posterior part of the ACC, which in turn connects with higher-order visual areas. This may suggest that the influence of area 47m on sensory processing is rather indirect and requires the mediation of other cortical regions (see [link](https://immersive.erc.monash.edu/neuromarvl/?save=45220d30-ce3b-4e17-922c-88964a883beb_31.16.56.20)).

#### Cingulate connectivities

In addition, there are further cycle-related changes in connectivity between the posterior part of the ACC and the higher-order ventromedial visual areas (VMV). Subregions along the ACC have been associated with salient sensory stimulation ^32,34^. The finding may represent the overly salient processing of sensory information^35^ throughout the migraine cycle due to the lack of habituation ^36^.

#### Normalisation of functional connectivity during the headache attack

We and others have argued previously that the underlying cause of migraine is an impairment to process the overflow of sensory information, which is ultimately reset through the headache attack. The overflow through the sensory stream that could not be contained ^5^ would make sensory information more salient which prevents habituation and may explain the current findings of the involvement of a number of bilateral insular subregions and regions in the posterior part of the anterior cingulate cortex. An ultimate decoupling of the connection between these regions may either contribute to the initiation of headache attacks in migraine or an active “reset” of cortical processes during the headache attack.

Taken together, the strong involvement of the regions aAIC and 47m in sensory and working memory processes suggest the current view on migraine as an overflow of sensory input that can not be sufficiently emptied from cortical loops that maintain conscious processes. This is in line with neurophysiological findings that interictal migraine patients suffer from the inability to habituate to sensory stimulation in the interictal phase ^3,36^. Similar to the connectivity of the aAIC and the BA47, habituation in migraineurs is reset to normal during and through a migraine headache attack ^3^. Future research is needed to address this hypothesis by directly probing salience and habituation throughout the migraine cycle.

#### A potential dysfunction of the agranular system

Interestingly, the AAIC and the posterior part of the orbitofrontal cortex are similar regarding gross morphology and architecture ^37^; they lack a granular layer 4 and are characterised by a substantial number of von Economo neurons (VENs ^26,38^). In their original work, von Economo and Koskinas (1925) stated that VENs can be found in “cap, corners and interior wall of the frontal part of the Gyrus limbicus, in the posterior transitional gyrus of the orbital part of the frontal lobe (which runs to the anterior frontoorbital Insula), in the so-called Gyrus transversus insulae and its associated Gyri breves accessorii (anteriores) insulae (translated by H.L. Seldon: [link](http://self.gutenberg.org/Get956uFile.aspx?&bookid=3468625))”^39^. A dysfunction of VENs may represent the underlying mechanism of the progressing impairment in episodic migraine ^40^; this needs to be specifically investigated with more appropriate techniques at the molecular level.

### Circularity of migraine related effects - potential limitations

One presupposition of our study is that we consider the migraine-phase-related processes as circular. In line with most studies, we assume the data from only one time point of data collection can be generalised to all potential occurrences of these circular time points. A clock metaphor may underline our assumptions on circularity;we tested our patients at 12 o’clock (the headache attack) and ended our day-by-day recording series at 11 o’clock (the prodrome phase), which is just before the next headache attack at 12 o’clock. We assume that a recording at 2 o’clock or any other ‘time’ would not be different across subsequent clock cycles or migraine cycles. Therefore, an attack recording (at 12 o’clock) should be representative for all attack recordings. Our analysis should not be considered as linear progression but as cyclic progression, which always restarts at 12 o’clock. This is the basis of interpretation of all studies on episodic migraine that compare the attack phase with a time point in the interictal phase. Although previous studies on interictal processes in migraine do not test circularity directly, the underlying assumption is the same, which is a differential effect of “time” during the cycle on cortical processing.

### Summary and Outlook

Most connectivity changes exhibited a linear increase over the pain-free interval with a peak prior to the headache and a “drop” during the headache (“reset mode”). Increasing functional connectivity towards the next attack is suggested to reflect the pathological sensory processing shown in previous behavioural and electrophysiological studies. The collapse of connectivity during the headache can be regarded as a mechanism of normalising cortical processing. Depending on the affected connection, the peak of synchronicity of functional connections during the ictal phase of the migraine cycle may contribute to the variety of symptoms during the headache phase of migraine attacks, such as headache, hypersensitivity to sensory modalities, and autonomous symptoms. These pathological processes are hypothesised to be related to VENs in agranular anterior insular and orbitofrontal brain regions. Future studies using more appropriate molecular techniques are needed to follow up on this hypothesis.

## Supporting information

see Supplementary Materials

## Data availability

Raw data were generated at the Technische Universität München. The authors confirm that the data supporting the findings of this study are available within the article. Inquiries for additional data are available from the corresponding authors, upon reasonable request.

## Acknowledgements

We thank Dr Andreas Straube for his comments and suggestions and Dr Stephanie Irving for copy-editing the manuscript.

