## Supplementary Table 1 for "Increased Functional Connectivity Characterises the Prodromal Phase of the Migraine Cycle"

**Tables**

**Supplementary Table 1.** Demographic characteristics and clinical migraine features of the measured and following headache attack.

| Patient | Age (years) | Female (f)/ male (m) | Attacks per month | Disease duration (years) | Measured headache attack | | | | | | Following headache attack | | | | | |
|---|---|---|---|---|---|---|---|---|---|---|---|---|---|---|---|---|
| | | | | | Attack severity (0-10) | Location of headache | With visual aura | Photo phobia | Phono phobia | Nausea (N)/ vomiting (V) | Attack severity (0-10) | Location of headache | With visual aura | Photo phobia | Phono phobia | Nausea (N)/ vomiting (V) |
| **1** | 28 | f | 3-6 | 6 | 6 | right-sided | yes | yes | yes | yes (N) | 6 | right-sided | yes | yes | yes | yes (N) |
| **2** | 24 | m | 3-6 | 14 | 5-6 | right-sided | no | yes | yes | no | 5 | right-sided | no | yes | yes | no |
| **3** | 26 | f | 3-6 | 24 | 6 | left-sided | no | yes | no | no | 6 | right-sided | no | yes | no | no |
| **4** | 40 | f | 1-2 | 29 | 7-8 | right-sided | yes | yes | yes | yes (N) | 6 | right-sided | yes | yes | yes | yes (N) |
| **5** | 32 | f | 1-2 | 15 | 6-7 | right-sided | no | yes | yes | no | 6 | right-sided | no | no | yes | no |
| **6** | 30 | f | 1-2 | 11 | 7 | bilateral | no | yes | no | no | 7 | bilateral | no | yes | no | no |
| **7** | 22 | f | 1-2 | 6 | 5 | left-sided | no | yes | yes | no | 6 | left-sided | no | yes | no | no |
| **8** | 23 | f | 3-6 | 7 | 8 | bilateral | no | yes | yes | yes (N) | 7 | right-sided | no | yes | yes | yes (N) |
| **9** | 33 | f | 6-10 | 23 | 8 | right-sided | no | yes | no | yes (N) | 8 | right-sided | no | yes | yes | yes (N, V) |
| **10** | 26 | f | 1-2 | 7 | 5-7 | left-sided | no | yes | no | no | 6 | left-sided | no | yes | no | no |
| **11** | 21 | f | 3-6 | 7 | 8 | bilateral | yes | yes | yes | no | 7 | bilateral | yes | yes | yes | no |
| **12** | 30 | f | 1-2 | 14 | 7 | bilateral | no | yes | yes | yes (N) | 7 | bilateral | yes | yes | yes | yes (N) |

Attack severity was recorded on a numerical rating scale ranging from 0 (no pain) to 10 (highest imaginable pain).

**Supplementary Table 2.** Migraine cycle related statistical significant connections for trajectory 1

| ROI 1 | ROI 2 | t-value | mean | std |
|---|---|---|---|---|
| Lateral Belt Complex | Frontal Opercular Area 1 | 6.67 | 0.20 | 0.07 |
| PreCuneus Visual Area | Frontal Opercular Area 3 | 6.31 | 0.33 | 0.18 |
| Insular Granular Complex | Area OP2-3/VS | 6.24 | 0.22 | 0.10 |
| Area p32 prime | Third Visual Area | 6.03 | 0.21 | -0.01 |
| VentroMedial Visual Area 2 | Anterior 24 prime | 5.96 | 0.19 | 0.06 |
| Area 47m | Area p32 prime | 5.95 | 0.20 | -0.01 |
| Auditory 5 Complex | Superior Frontal Language Area | 5.87 | 0.24 | 0.06 |
| Area 47m | RetroInsular Cortex | 5.87 | 0.21 | 0.13 |
| cerebellum_24_Vermis_IX | Area anterior 9-46v | 5.83 | 0.21 | 0.36 |
| Frontal OPercular Area 1 | Primary Auditory Cortex | 5.81 | 0.18 | 0.04 |
| Area 47m | Area p32 prime | 5.76 | 0.33 | 0.02 |
| CEREBELLUM_RIGHT | Fusiform Face Complex | 5.74 | 0.20 | 0.07 |
| Area 47m | Area 8C | 5.73 | 0.23 | 0.36 |
| Area OP1/SII | Area 31p ventral | 5.70 | 0.21 | 0.15 |
| Anterior Agranular Insula Complex | Third Visual Area | 5.69 | 0.15 | 0.07 |
| Entorhinal Cortex | PeriSylvian Language Area | 5.69 | 0.20 | 0.14 |
| Area 47s | Ventral Area 6 | 5.68 | 0.22 | 0.14 |
| Lateral Belt Complex | Anterior Agranular Insula Complex | 5.65 | 0.17 | 0.02 |
| Frontal OPercular Area 1 | Area 52 | 5.62 | 0.17 | -0.04 |
| ProStriate Area | Area anterior 32 prime | 5.61 | 0.18 | 0.05 |
| CEREBELLUM_RIGHT | Eighth Visual Area | 5.57 | 0.19 | 0.00 |
| Area 47m | Premotor Eye Field | 5.55 | 0.26 | -0.21 |
| Frontal OPercular Area 2 | Area OP2-3/VS | 5.54 | 0.20 | 0.08 |
| ProStriate Area | Anterior 24 prime | 5.51 | 0.21 | 0.09 |
| Anterior Agranular Insula Complex | Area V4t | 5.50 | 0.23 | 0.11 |
| Area Frontal Opercular 5 | Area posterior 9-46v | 5.49 | 0.21 | 0.15 |
| VentroMedial Visual Area 1 | Anterior 24 prime | 5.48 | 0.29 | 0.07 |
| Entorhinal Cortex | Premotor Eye Field | 5.45 | 0.19 | 0.02 |
| Entorhinal Cortex | Premotor Eye Field | 5.45 | 0.22 | -0.09 |
| Area V6A | Anterior Agranular Insula Complex | 5.44 | 0.20 | 0.10 |
| Area 47m | Frontal OPercular Area 3 | 5.42 | 0.21 | -0.02 |
| THALAMUS_RIGHT | PosteriorInsular Area2 | 5.41 | 0.29 | 0.16 |
| Area 47m | Supplementary and Cingulate Eye Field | 5.37 | 0.21 | -0.03 |
| Area 47m | Frontal Eye Fields | 5.35 | 0.22 | 0.27 |
| Anterior Agranular Insula Complex | Superior Temporal Visual Area | 5.34 | 0.23 | -0.02 |
| Area 47s | Area p32 prime | 5.33 | 0.19 | 0.16 |
| Area OP4/PV | Area 47s | 5.32 | 0.21 | 0.08 |
| PALLIDUM_RIGHT | Frontal OPercular Area 3 | 5.32 | 0.18 | 0.27 |
| cerebellum_01_Left_I-IV | Area posterior 9-46v | 5.32 | 0.22 | 0.01 |
| Anterior Agranular Insula Complex | AreaTemporoParietoOccipital Junction 2 | 5.31 | 0.17 | 0.05 |

| | | | | |
|---|---|---|---|---|
| cerebellum_28_Right_X | cerebellum_03_Left_V | 5.31 | 0.14 | 0.08 |
| AreaPosteriorInsular1 | Area 3a | 5.31 | 0.24 | 0.00 |
| VentroMedial Visual Area 1 | Area anterior 32 prime | 5.30 | 0.16 | 0.19 |
| THALAMUS_LEFT | PALLIDUM_RIGHT | 5.28 | 0.24 | -0.11 |
| Insular Granular Complex | Area 31p ventral | 5.28 | 0.23 | 0.04 |
| Area IntraParietal 0 | Area 46 | 5.27 | 0.22 | 0.11 |
| Area 47m | Area OP2-3/VS | 5.27 | 0.17 | 0.27 |
| Primary Auditory Cortex | Frontal OPercular Area 3 | 5.26 | 0.16 | -0.02 |
| Area OP1/SII | Area 31p ventral | 5.26 | 0.20 | 0.11 |
| Anterior Agranular Insula Complex | Area V3A | 5.25 | 0.20 | 0.21 |
| Anterior Agranular Insula Complex | Area Lateral Occipital2 | 5.25 | 0.20 | -0.03 |
| Area 31pd | Area PFcm | 5.25 | 0.18 | 0.14 |
| Area p32 prime | Area V3A | 5.24 | 0.21 | 0.09 |
| cerebellum_01_Left_I-IV | Entorhinal Cortex | 5.24 | 0.26 | -0.08 |
| Area 47m | Area 44 | 5.23 | 0.20 | 0.07 |
| Anterior Agranular Insula Complex | Primary Auditory Cortex | 5.23 | 0.33 | 0.18 |
| Area p32 prime | Second Visual Area | 5.23 | 0.22 | 0.10 |
| MiddleTemporalArea | Area posterior 24 | 5.23 | 0.21 | -0.01 |
| Area 47m | Primary Motor Cortex | 5.22 | 0.19 | 0.06 |
| Area p32 prime | PreCuneus Visual Area | 5.21 | 0.20 | -0.01 |
| VentroMedial Visual Area 1 | Area 8BM | 5.21 | 0.24 | 0.06 |
| Area anterior 32 prime | Area 47m | 5.20 | 0.21 | 0.13 |
| Anterior Agranular Insula Complex | Area 55b | 5.20 | 0.21 | 0.36 |
| Area PF opercular | Area 31pd | 5.20 | 0.18 | 0.04 |
