## Supplementary material for "Increased Functional Connectivity Characterises the Prodromal Phase of the Migraine Cycle": see Supplementary Materials

### Challenges of assessing circular processes in episodic migraine

*Q: How did the authors adjust the data based in terms of movement inside the scanner, anxiety, depression, BMI, hypertension, physical exercise level, smoking status, alcohol use, sex, headache frequency, number of total scans and analgesics use?*
A: There is a plethora of factors that may influence the cortical processing of migraine. A proper control of the proposed factors would require a much larger number of subjects equally represented for the different levels of the proposed factors (e.g. equal or comparable number of men and women, a comparable number of subjects for different headache frequency groups, etc.). Each added explanatory factor must be fed by additional data. Adding all of these factors into the longitudinal analysis would require hundreds of recordings. Nevertheless, the above results have the potential to answer the basic question stated in the study, which is to investigate the influence of the time point of the migraine cycle on cortical processing. A recent study [2] has challenged this type of analysis for limited sample sizes.

*Q: How does the parcellation work? Did the authors perform a ROI-to-ROI analysis or on the contrary a cluster-wise analysis? Can the data be presented as a 3D brain image?*
A: The analysis is basically a multiple region-of-interest (ROI) analysis. The boundaries of the regions are defined by the Glasser Atlas [1]. We did not do any statistics on single voxels but ran a PCA on the entire ROI and extracted the first component. For this reason, there is no maximum *t*-value with a specific MNI coordinate available. The individual ROIs were defined by individual 2D surface projections. For this reason there is no common MNI coordinate.

*Q: Why did the authors do a spatial smoothing of 5 mm?*
A: We were also interested in subcortical regions; small nuclei often do not exceed a diameter of 5mm.

Q: *Does the randomisation approach to determine the significance threshold also correct for multiple comparisons?*
A: Yes.

*Q: The length of the cycle differs and the number of recordings for each subject, too? Is there a way to deal with this?*
A: In an ideal case, we would have acquired the same number of recordings from patients with an identical cycle length. However, the length of the migraine cycle is genuinely variable and largely unpredictable. This means that we had to inevitably deal with either different numbers of recordings or variable gaps between recordings. We decided to have a reasonable compromise between both extrema.
For estimating the slope of the regression line, the first and last data points of the cycle have the strongest impact and these are the time points of the migraine cycle we were mostly interested in. Consequently, the follow-up recording after the headache attack started the next day after the headache attack. For all patients, we collected the last data 48h before the following headache attack. 10 out of the 12 patients were recorded one day before the following headache attack. For economical reasons, the other interictal

recordings were evenly distributed along the cycle with gaps between 1 day (short cyclers) and 4 days (long cyclers). The statistical model does not require the same number of recordings or the same cycle length. We collected sufficient data for fitting the regression parameters. See figure from our previous publication on the same dataset.

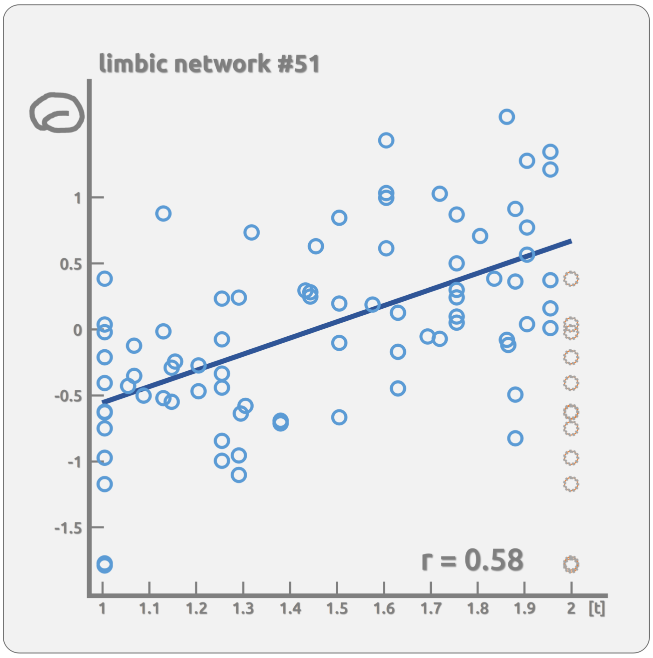


The x-axis shows the normalised time course of the migraine cycle. Time point 1 and 2 represent the headache attack days.

*Q: Does the model account for the different number of recordings?*

It is important to note that mixed linear models can handle unbalanced data very well. Thus, the uneven representation of cases with different frequencies of headaches is not a problem in terms of the quality of the estimated fixed factors (model parameters). Therefore, a linear mixed-effects model is a suitable tool for modelling unbalanced data. In addition, the number of recordings could be included in the model under the condition that there would be a systematic impact on the findings. For example, longer migraine cycles result in a higher number of recordings than shorter cycles. However, the idea that there might be systematic differences in the pathophysiology of short vs. long cyclers does not appear plausible.

*Q: Why do the authors call the time courses “trajectories"?*

A: The term trajectory is certainly a metaphor that is used in medicine and developmental science to describe the progress of language or cognitive capabilities. The term “trajectory” underlines the determined and cyclic nature of the migraine attacks. A simple “time course 1” or “time course 2” would sound too passive and the semantics would suggest processes that just somehow happen. Therefore, in our view, the term trajectory is still best suited to describe *predefined* and finally inevitable time courses of neuronal activity.

The shape of the trajectory is predefined and other shapes would be potentially plausible. For the present study we decided to assume a linear progression towards the next attack (or the day before). However, a non-linear quadratic or any other exponential progress might be plausible, too. There might be some differences in taste regarding the use of metaphoric terminology. Metaphoric terminology with temporal connotations such as “evolve”, “emerge” and “unfold” clearly help to make scientific work more digestible.

*Q: Why is there no control group included?*
A: This is not possible for the present study. The reason is that there is nothing comparable to the migraine cycle. For the patients we encoded the migraine cycle with numbers from 1 to 2. For example, for a recording exactly in between two attacks we assigned the number 1.5 to this respective recording as we assumed that the magnitude of brain activity in the middle of the cycle would be halfway between the first and the second attack. This principle does not work for any control group, where we would need to assume the same "number" for each recording. The fixed effect of a LME would naturally be a null result. This is comparable with a correlation between two time series: 1 to 10 correlated with ten times the number 5 for which r=NaN. [in Matlab : correlation = corr((1:10)',5+zeros(10,1))]

*Q. How can you assume a change of connectivity for the second headache phase if participants were not scanned during a second migraine?*

We are interpreting our results as circular and not as linear processes. Based on previous neurophysiological research, we assume that the processes that we describe in our analysis will occur repeatedly in a very similar fashion. Otherwise, circular processes (e.g. in chronobiology) would always need a certain number of repetitions. This also applies to all studies on migraine that specifically investigate the selected phases of the migraine cycle. Most studies include just one recording per patient and cycle point; they assume that ONE recording of a subject represents ALL potential recordings of that subject.
In line with the current interpretation of neuroimaging findings, we can indeed assume that ONE postdrome recording reflects ALL postdrome recordings. Challenging this assumption would imply that the findings of ANY study would need a second or multiple confirmations of the same mental or physical state of a subject. Studies investigating cortical processes during the ictal phase compared to the headache phase are assumed to be valid in the follow-up migraine cycle. Although these studies do not test circularity, the underlying assumption is the same, which is that one recording represents all potential follow-up recordings.

*Q: The imaging time series for each patient started with the recording of a spontaneous, untriggered and untreated headache attack within the first 6 hours after the beginning of the headache. Does this mean that the first scan performed in each patient was during a migraine attack*?
A: Yes.

*Q: How has the statistical threshold been determined and has the statistics been corrected for multiple testing?*
A: Using randomised time vector data, the entire LME analysis was repeated 5000 times, resulting in 5000*89675 statistical tests. The highest absolute t-values of each of the 5000

repetitions were extracted. This procedure resulted in a right-skewed distribution of 5000 values. Based on this distribution, the statistical threshold was determined using the “palm_datapval.m” function publicly available in PALM [3,4]. A test was considered as “significant” if it exceeded the threshold provided by PALM ($p<0.05$), which is equivalent to a t-value of 5.195 of the original test.

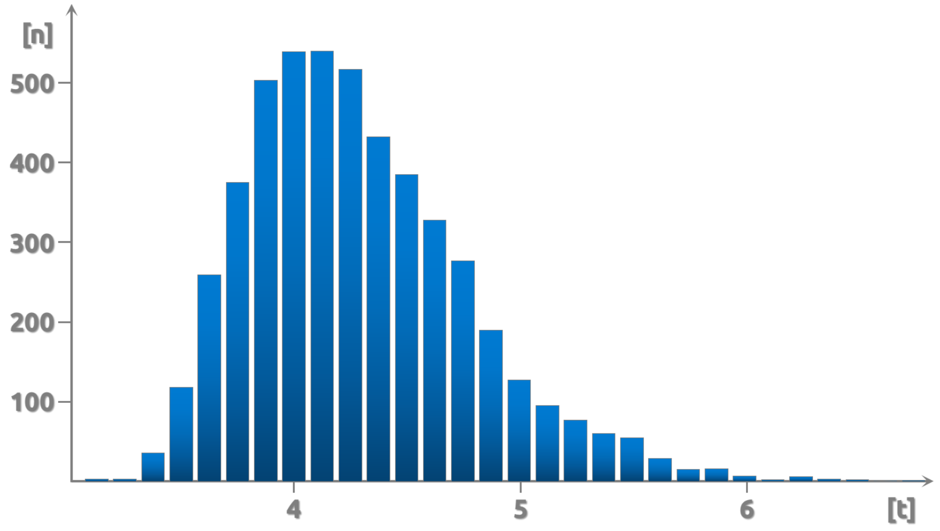


**Distribution of statistics on randomised data.** The distributions show the absolute maximal values for each of the 5000 randomisations.
